# Autophagy impairment by ATG4B deficiency reduces experimental hypersensitivity pneumonitis severity

**DOI:** 10.64898/2026.03.16.712004

**Authors:** Sandra Cabrera, Andrea Sánchez, Miguel Gaxiola, Ángeles García-Vicente, Moisés Selman, Annie Pardo

## Abstract

Autophagy has been implicated in several lung diseases, either protecting tissues or driving pathology. Hypersensitivity pneumonitis (HP) is a complex inflammatory lung disease, and autophagy is heavily involved in regulating inflammation. The role of autophagy in HP remains unclear. The aim of our study was to understand the role of autophagy in HP pathogenesis. GFP-LC3 transgenic mice were exposed intranasally to *Saccharopolyspora rectivirgula* (SR) to induce HP and follow autophagy activation in the lung. Then, we take advantage of our Atg4b-deficient mouse model to assess how autophagy disruption impacts lung inflammation in response to SR antigen challenge. Increased autophagy activation was observed in epithelial and inflammatory cells after SR antigen exposure in GFP-LC3 transgenic lungs. GFP-LC3 puncta colocalized with ATG4B and ATG5 in epithelial and inflammatory cells after antigen exposure. Autophagy impairment limits the inflammatory response after SR antigen exposure in the lungs from the *Atg4b*-deficient mice when compared to WT mice. To evaluate whether lipopolysaccharide (LPS) exacerbates the inflammatory response in the Atg4b-deficient, a SR+LPS combined treatment was developed and we discovered that LPS aggravates the SR-induced HP in WT but not in Atg4b-deficient mice. Reduced HP severity in *Atg4b*-deficient mice was associated with decreased expression of NFkB, CCL1, CCL25, CXCL1, TNFR1, IL-13, and IL-17A, diminished CD4+ T cell recruitment and expansion, reduced M2-like macrophages, and decreased granuloma and iBALT development. Our findings highlight autophagy as a critical driver in HP pathogenesis and as a potetial target for novel theraphy development.

## INTRODUCTION

Hypersensitivity pneumonitis (HP) is an interstitial lung disease (ILD) caused by repeated exposure to inhaled antigens in genetically predisposed individuals [1–3]. Those antigens may be organic, for example microorganisms (bacteria and fungi), and proteins derived from plants and animals, but also inorganic agents. The most frequent antigens that trigger HP are derived from bird proteins (Bird Fancier’s/ Pigeon breeders’ disease) and from the actinobacteria *Saccharopolyspora rectivirgula* (Farmer’s Lung disease). Chronic antigen exposure can occur indoors or outdoors at work, home or recreational environments [3–5].

The disease is categorized into non-fibrotic and fibrotic forms [3–5]. Non-fibrotic HP is characterized by inflammation mediated by alveolar macrophages, dendritic cells, and T helper cell type 1 (Th1) and T helper cell type 17 (Th17) cells. Fibrotic HP is characterized by persistent inflammation, directed by a shift from Th1 to Th2 and persistent Th17 cells, that leads to fibroblast activation, extracellular matrix (ECM) deposition, and lung fibrosis [4, 5].

Autophagy is a cellular degradation process that removes damaged organelles and proteins, and is essential for lung proteostasis and homeostasis. Despite the protective role of autophagy on cells, abnormal or excessive autophagic responses cause several pathological conditions [6]. Autophagy plays both protective and detrimental roles in lung diseases; for example, in idiopathic pulmonary fibrosis (IPF) it can be protective, and its dysregulation promotes fibrosis, while in asthma and chronic obstructive pulmonary disease (COPD) excessive autophagy can contribute to epithelial apoptosis, lung damage, and chronic inflammation [7,8].

The contribution of autophagy in HP pathogenesis remains unclear. We have recently shown increased expression of key autophagic proteins like LC3B, p62, ATG4B, and ATG5 in epithelial and inflammatory cells in the lungs of HP patients compared to healthy individuals, suggesting autophagy could be involved in HP pathogenesis [9, 10]. The aim of this study was to investigate if autophagy has beneficial or detrimental effects after *Saccharopolyspora rectivirgula-induced*

experimental HP using our *Atg4b* knockout mouse model that shows a systemic severe reduction in autophagy activity [11]. Understanding the effects of autophagy on disease pathogenesis and progression may potentially contribute to designing novel strategies for HP treatment.

## METHODS

### Mice and *Saccharopolyspora rectivirgula*-induced experimental HP model

The C57BL/6J, *Atg4b*^−/−^ and GFP-LC3 transgenic mice were maintained under specific pathogen-free (SPF) conditions. The generation of *Atg4b*−/− and GFP-LC3 transgenic mice have been previously described [11–13]. The mice used were 8-10 weeks of age at the start of each experiment. The experiment was approved by The Bioethics Committee of the Faculty of Sciences from Universidad Nacional Autónoma de México (protocol approval PI_2022_23_07_Cabrera) and the National Institute of Respiratory Diseases of Mexico (B24-23) and performed in strict accordance with approved guidelines.

Mice were exposed intranasally with *Saccharopolyspora rectivirgula* (*SR* (ATCC 15347)) 50 μg / mouse, alone or in combination with 5μg of Lipopolysaccharide (LPS from *Escherichia coli* O111:B4 (sc-3535B)), or sterile saline 3 times per week for 3 weeks, and were euthanized 3 days after the last exposure. Mice were sedated with isoflurane to allow intranasal inhalation of either sterile saline, SR, or SR+LPS (50μl).

### Morphology

Mice were perfused transcardially with sterile saline, and lungs were inflated with 4% neutral-buffered formalin at a continuous pressure of 25 cm H₂O and fixed. Paraffin-embedded sections (5 μm) were stained with hematoxylin and eosin (H&E) and assessed semi-quantitatively for inflammation, severity of lung damage, and extent of lung lesions. The injury score represents: 0 score - no lung injury; 1 - mild lung injury; 2- moderate lung injury; and 3- severe lung injury.

### Immunohistochemistry

The tissue sections were deparaffinized and were then rehydrated and blocked with 3% H2O2 in methanol, followed by antigen retrieval in a microwave in 10 mM citrate buffer, pH 6.0. Tissue sections were treated with universal blocking solution (BioGenex, HK085–5K) for 10 min, and then incubated overnight at 4°C with the following primary antibodies: LC3B (L7543), anti-ATG4B (A2981), anti-CD4 (SAB5500064) and anti-ARG1 (SC-271430) all from Sigma- Aldrich, anti-RELA/NFκB p65 (Abcam, ab7970), and anti-YM1 (Stem Cell Technologies, 1404). A secondary biotinylated anti-immunoglobulin followed by horseradish peroxidase-conjugated streptavidin (BioGenex, HK330–5K) was used according to the manufacturer’s instructions. 3-amino-9-ethyl-carbazole (BioGenex, HK092–5K) in acetate buffer containing 0.05% H2O2 was used as the substrate. The sections were counterstained with hematoxylin. The primary antibody was replaced by nonimmune serum for negative control slides.

### Imaging acqusition and analysis

Lung images were taken using 10X, 20X or 40X objectives from the Eclipse E600 Nikon microscope (Nikon, New York, USA) equipped with a DXM1200C digital camera (Nikon, New York, USA) and the Nis Elements 3.0 software (Nikon, New York, USA). From each slide, 12 nonoverlapping fields per mouse were captured at 40X and analyzed with the QuPath bioimage analysis software (https://qupath.github.io). A full-image annotation was created from the brightfield digital images and the QuPath’s Positive Cell Detection algorithm was performed (Percentage of positive cells = positive cells / total nuclei × 100) [14]. The mean percentage of positive cells from all fields was calculated for each animal and plot. The total tissue area that represents the entire lung sample in the slide and the inducible bronchus-associated lymphoid tissue (iBALT) area (µm^2^) were calculated by using the free-hand tool to draw these regions with ImageJ/FIJI software.

### Immunofluorescence Analysis

Lungs from GFP-LC3 transgenic mice were analyzed with a fluorescence microscope; dots were observed and counted in 3 independent visual fields from at least 6 independent mice. The acquired images were deconvolved using LAS X software to improve resolution, enabling higher-quality visualization of dots. For colocalization, tissue slides were processed equally as IHC slides and incubated with the following primary antibodies: anti-ATG4B (A29981), anti-ATG5 (A0731), anti-SPC (AB3786) and anti-CD4 (SAB5500064) from Sigma-Aldrich, anti-CD68 (MU549-UC), anti-MPO (PU496-5UP), and anti-CD19 (MU231- 5UC) from Biogenex, and anti-Krt8 (Thermo Fisher Scientific, MA5-32118). After washing, tissues were incubated with secondary antibodies Alexa 647 anti-rabbit (A31573), Alexa 546 anti-mouse (A11030) and Alexa 546 anti-Rabbit (A-11035) from Thermo Fisher Scientific. Nuclei were stained with DAPI (Blue) (D1306, Thermo Fisher Scientific). Samples were imaged with a Confocal Laser Scanning Microscope TCS SP8 using a CS2 Plan Apochromat 63× (Leica, Wetzlar, Germany). Background noise for the fluorophore emitting at 546, 647, and DAPI channels was reduced by subtracting the negative noise signal using the Fiji program and applied to the entire images. A minimum of six representative, non-overlapping fields from lungs were evaluated per tissue section at 63X magnification. Regions of interest were cropped and enlarged to show greater detail in digital insets.

### Bronchoalveolar lavage fluid (BALF) analysis

Mice were intubated with a 20G ½ catheter (BD insyte autoguard, 381834) and lungs were lavaged with 1.0 mL of sterile saline. Cells were isolated from BALF by centrifugation at 2,000 × g for 10 min at 4°C, and the supernatant was stored at - 80°C for cytokine analysis. Total cell counts were performed under light microscopy with a standard hemocytometer and differential cell counts were obtained using Wright-Giemsa stain and performed by the pathologist.

### Mouse cytokine antibody array

Mouse inflammation antibody arrays (ab133999) were used to quantify in BALF samples from either controls or SR+LPS-exposed WT and *Atg4b*^-/-^ mice, and were performed according to the manufacturer’s instructions. A total of 40 cytokines were included in the array as follows: BLC (CXCL13), CD30 ligand, CCL1, MPIF-27CCL24, TNFSF6, CX3CL1, G-CSF, GM-CSF, IFN 𝝲, IL-1𝝰, IL-𝝱, IL-2, IL-3, IL-4, IL-6, IL-9, IL-10, IL-12 p40/ p70, IL-12 p70, IL-13, IL-17A, CXCL11, CXCL1, Leptin, LIX, XCL1, CCL2, M-CSF, CXCL9, CCL3, MIP-1, CCL5, CXCL12 𝝰, CCL1, CCL25, TIMP-1, TIMP-2, TNF-𝝰, TNF RI and TNF RII. Array images were captured, and spot intensities were quantified using ImageJ. Relative cytokine levels were compared after densitometry analysis. Results are expressed as fold change.

### Statistics

All experimental data are reported as mean ± SEM. Datasets were tested for normality and homogeneity of variances. Comparisons between two groups were performed using Student’s t-test. In experiments with more than 2 groups, differences were analyzed using a multifactorial one- or two-way analysis of variance (ANOVA) followed by a Tukey post hoc test. All statistical analyses and graphical representations were performed using GraphPad Prism software Version 10.6.1 (Graphpad Software Inc., San Diego CA) and *P* values lower than 0.05 were considered significant.

## RESULTS

### Autophagy is increased in epithelial and inflammatory cells in the lung after *Saccharopolyspora rectivirgula* exposure

To investigate whether autophagy is induced after *SR* challenge, C57BL/6 mice were exposed intranasally to *SR* antigen 3 days a week for 3 weeks. Histologic analyses of *SR*-exposed lungs revealed increased perivascular, peribronchiolar, intraalveolar and interstitial inflammation compared to controls. A low positive staining for the autophagy biomarker LC3B was observed in some macrophages in control lungs, while a strong LC3B-positive staining was observed in inflammatory and epithelial cells in the *SR*-exposed HP lungs (**Fig. 1a**). LC3B-positive cells were significantly increased in *SR*-exposed lungs compared to controls (**Fig. 1b**). The cysteine-protease ATG4B, essential for the pro-LC3B maturation [11, 15], was found significantly increased in bronchial epithelial and inflammatory cells in HP lungs, while no signal was observed in control lungs (**Fig. 1a** and **c**).

**Figure 1.**
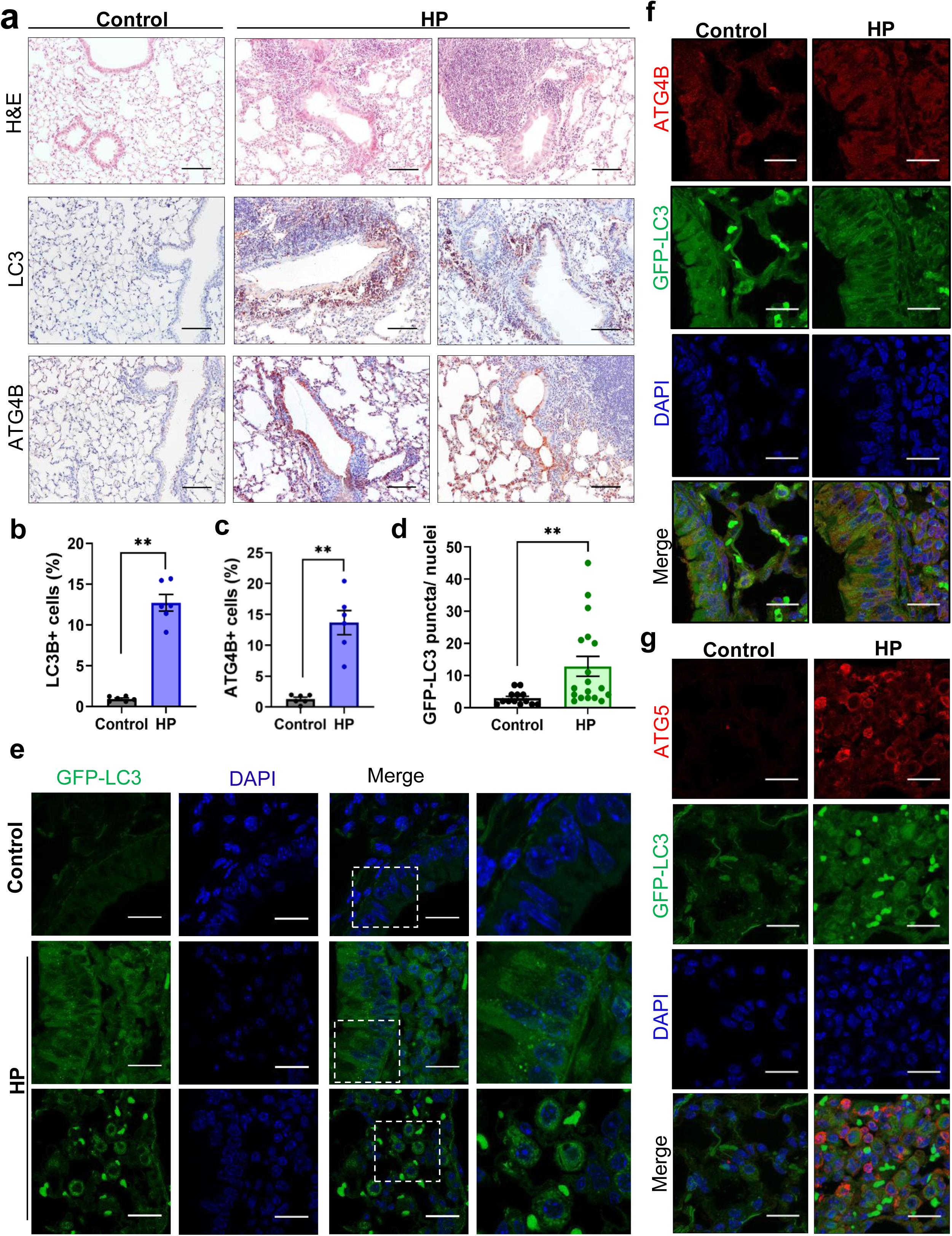
Autophagy induction after experimental HP. **a**. Representative light microscopy images of H&E-stained lung tissue sections from control and experimental HP mice (Images are at 10X, and scale bars represent 20 µm). Representative photomicrographs of immunohistochemical staining for LC3B and ATG4B in controls and *Saccharopolyspora rectivirgula (SR)*-exposed mice. Positive staining is observed in red, and nuclei are counterstained with hematoxylin in blue. Images are at 20X magnification (Scale bars = 100). **b.** Percentage of LC3B-positive and **c.** ATG4B positive cells in control and HP lungs. **d**. Quantification of LC3 puncta per nuclei of lung tissue from control and HP-lungs from GFP-LC3 transgenic mice. **e.** Representative fluorescent microphotographs of control and HP-lungs from GFP-LC3 transgenic mice. GFP-LC3 puncta are observed in green and DAPI-stained nuclei in blue. Images are at 63X (Scale bars = 20 µm), and the digital insets (white dashed box) represent a 5 times magnification. **f.** Representative fluorescent microphotographs of ATG4B and **g.** ATG5 colocalization with GFP-LC3 signal. Data are presented as mean ± SEM. Statistical significance was determined with t-test (**p<0.005).

Then, the GFP-LC3 transgenic mice were used to induce experimental HP and assess autophagosome formation. Control transgenic lungs showed a GFP-LC3 diffuse cytoplasmic signal, with some cells showing few GFP-LC3 cytoplasmic puncta, indicating a low LC3-lipidated form under non-stressed conditions. Increased GFP-LC3-positive dots per nucleus, indicating LC3B incorporation into autophagosome membranes, were significantly increased in *SR*-exposed HP transgenic lungs (**Fig. 1d** and **e**). Increased ATG4B colocalization with GFP-LC3 puncta in HP lungs from transgenic mice was observed mainly in bronchial epithelial cells and macrophages in HP when compared to control lungs (**Fig. 1f**). Increased positive signal for ATG5 (an E3-like ligase from the ATG12 conjugation system) [6, 7] and colocalization with GFP-LC3 puncta were observed in macrophages from HP compared to control transgenic lungs (**Fig. 1f**).

Afterwards, we used cell-specific antibodies to identify different cell subtypes undergoing autophagy after HP induction by *SR* exposure. We observed colocalization of GFP-LC3 puncta with keratin 8 (KRT8), a marker of simple columnar epithelial cells from airways, and surfactant C protein (SPC), a marker of alveolar type 2 epithelial cells in HP lungs from GFP-LC3 transgenic mice (**Fig. 2a**). We also identified CD68 positive cells, a pan-biomarker of macrophages, and myeloperoxidase (MPO) positive cells, a biomarker of neutrophils, to colocalize with GFP-LC3 puncta (**Fig. 2b**). Additionally, GFP-LC3 puncta colocalization with anti-CD19 and anti-CD4 positive signal, revealed that autophagy is activated also in B and T lymphocytes (**Fig. 2c**). In summary, our findings demonstrated that autophagy is activated in bronchial and alveolar epithelial cells, but also in innate and adaptive inflammatory cells after experimental HP.

**Figure 2.**
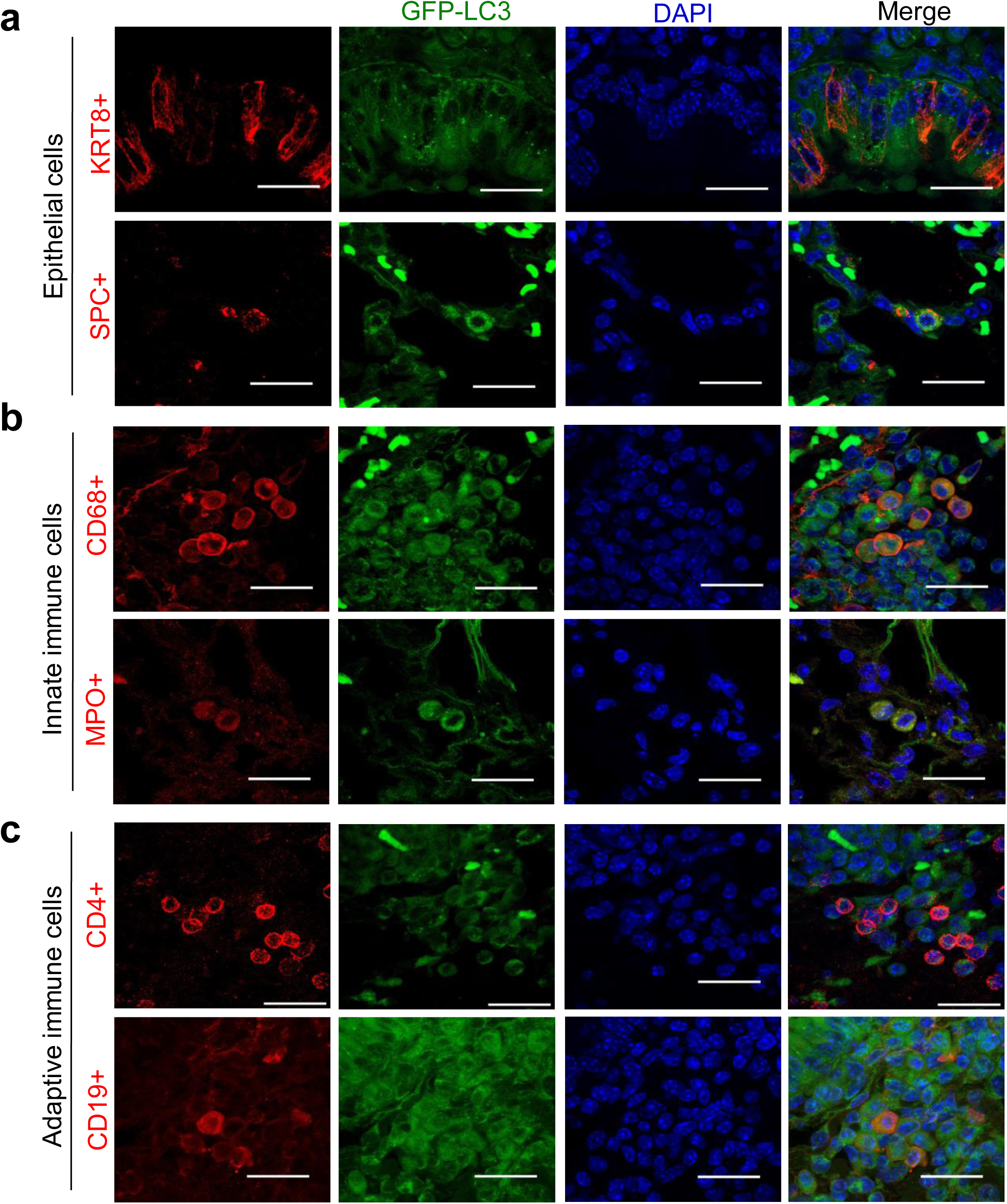
Epithelial and immune cell biomarker colocalization with GFP-LC3B positive puncta in HP lungs. **a.** Representative confocal microphotographs of keratin 8 (KRT8), a biomarker of simple columnar epithelial cells form airways, and surfactant C protein (SPC), a biomarker of alveolar type 2 epithelial cells in HP lungs from GFP-LC3 transgenic mice. **b.** Representative confocal microphotographs of CD68+ macrophages and myeloperoxidase (MPO) positive cells (a biomarker of neutrophils) in HP lungs from GFP-LC3 transgenic mice. **c.** Representative confocal microphotographs of CD19+ (B cells) and CD4+ (T cells) in HP lungs from GFP-LC3 transgenic mice. All biomarker-specific positive cells are in red, GFP-LC3B dots in green, and DAPI-stained nuclei are in blue. Images are at 63X (Scale bars = 20 µm).

### *Atg4b⁻/⁻* mice display an attenuated inflammatory response after *Saccharopolyspora rectivirgula* challenge

We have previously demonstrated that ATG4B absence inhibits autophagic activity in most tissues including the lung [11, 15]. In this study, *Atg4b*-deficient mice were used to induce experimental HP, with the aim of evaluating the role of autophagy in HP pathogenesis. Control lung tissue sections yielded no apparent morphological differences between WT and *Atg4b*^-/-^ control mice, as previously described (**Fig. 3a** and Supplementary Fig. 1). After 3 weeks of *SR* intranasal instillation, increased mononuclear cell infiltration around peribronchiolar and perivascular regions, intraalveolar inflammation, and thickening of the alveolar wall were observed in WT compared to *SR*-exposed *Atg4b*^-/-^ lungs (**Fig. 3a** and **b** and Supplementary Fig. 1). Since we observed a poor inflammatory response in *Atg4b*^-/-^ mice after *SR*, we decided to use a combination of the antigen with LPS as a second hit to exacerbate the inflammatory response. LPS single treatment produced thickening of the alveolar wall and increased intra-alveolar inflammation (mainly neutrophils and macrophages) in both WT and *Atg4b*^-/-^ mice compared to controls (**Fig. 3a** and **b**). *SR* challenge in combination with LPS increased the injury score in WT, but no in the *Atg4b^-/-^* mice. Moreover, LPS addition exacerbated the development of inducible bronchus-associated lymphoid tissue (iBALT) in 83% of the *SR*+LPS-exposed WT, while only 17% of *SR*+LPS-exposed *Atg4b^-/-^* showed iBALT development, and the iBALT area was significantly more prominent in WT compared to *Atg4b^-/-^* mice (**Fig. 3a** and **c**, and Supplementary Fig. 2). Moreover, *SR* exposure in combination with LPS elicited granuloma development in 50% of *SR*+LPS-exposed WT mice, while one granuloma was observed in the lung parenchyma from only one knockout mouse (17% of the *Atg4b*^-/-^ *SR*+LPS-exposed mice, Supplementary Fig. 2).

**Figure 3.**
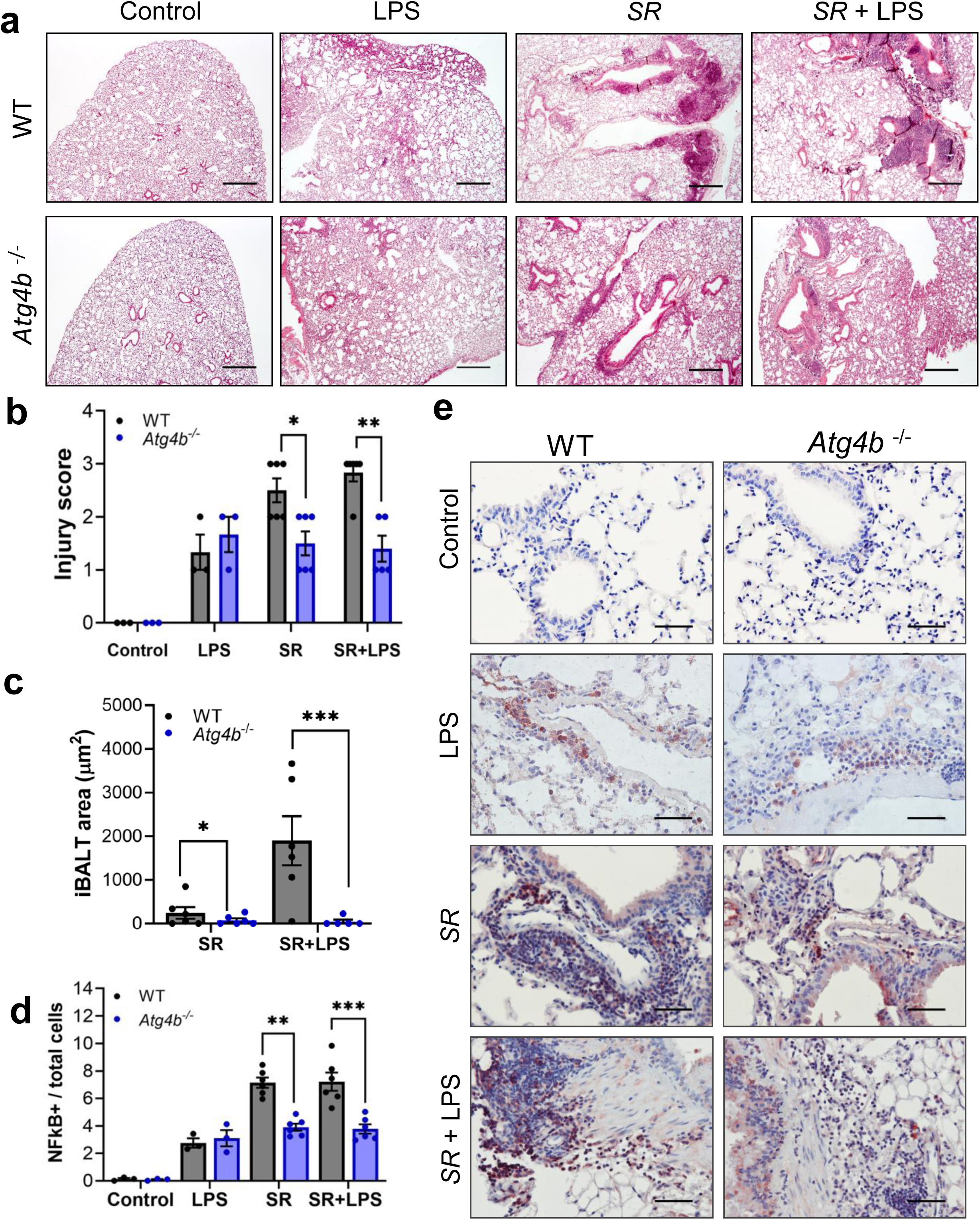
Reduced inflammatory response and HP severity in *Atg4b*^-/-^ mice. **a.** Representative light microscopy images of H&E-stained lung tissue sections from control, LPS-, SR- and SR+LPS-exposed WT and *Atg4b*^-/-^ mice after 3 weeks of treatment. Images are at 10X, and scale bars represent 200 µm. **b.** Injury score represents: 0 score - no lung injury; 1 - mild lung injury; 2- moderate lung injury and 3- severe lung injury. **c.** iBALT area quantification relative to whole lung lobe area from SR+LPS exposed WT and *Atg4b*^-/-^ mice. **d**. NFkB-positive cells per total nuclei (total cell) per experimental group. **e.** Representative photomicrographs of immunohistochemical positive staining for NFkB in control, LPS-, SR- and SR+LPS-exposed WT and *Atg4b*^-/-^ lungs. Images are at 20X magnification (Scale bars = 100).Data are presented as mean ± SEM. Statistical significance was determined by multifactorial two-way analysis of variance (ANOVA) followed by a Tukey post hoc test (* p<0.05, ** p<0.005, ***p<0.0001).

Total inflammatory cells in BALF were significantly increased in LPS, *SR* and *SR*+LPS challenged mice from both WT and *Atg4b*^-/-^ mice compared to their saline controls. Paradoxically, there was no difference in BALF total cell count between *Atg4b*^-/-^ mice and their WT littermates after LPS alone, *SR* alone or *SR* combined with LPS. BALF cell differential analysis revealed a marked heterogeneity in macrophages, lymphocytes and neutrophils in both WT and *Atg4b*^-/-^ mice exposed to LPS, *SR* and *SR*+LPS (Supplementary Fig. 2). Inflammatory cells in BALF did not accurately reflect the differences we observed in the lung parenchyma between *SR* and *SR*+LPS treated WT and *Atg4b*^-/-^ mice.

Then, we evaluated the localization of nuclear factor kappa B (NFkB) in lung tissue from WT and *Atg4b*^-/-^ mice, since it is a master transcription factor that drives the expression of proinflammatory cytokines and modulates inflammation. We found an increased and strong NFkB positive staining in bronchial epithelial cells, neutrophils and lymphocytes in lungs from both *SR* and *SR*+LPS-exposed WT compared to *Atg4b*^-/-^ exposed mice (**Fig. 3d** and **e**). Our data indicates that impaired autophagy, by the loss of ATG4B, significantly reduced inflammation in the lung parenchyma after SR-exposure, and even with the sum of LPS as a second insult.

### Reduced Th1, Th2 and Th17 cytokines in BALF, and lower CD4+ T-cells and M2-like macrophages in *Atg4b*^-/-^ lung parenchyma after experimental HP

To better understand the lung inflammatory milieu in the lungs from *Atg4b*^-/-^ and WT mice after experimentally induced HP, we performed a multiplex cytokine antibody array to analyze 40 cytokines in BALF samples. The heat map represents the fold change in protein expression relative to the WT control group. We did not find differences in the BALF cytokines derived from *Atg4b*^-/-^ and WT control lungs. After experimental HP induction, the following cytokines were significantly increased in the BALF derived from *SR*+LPS-exposed WT compared to *SR*+LPS-exposed *Atg4b*^-/-^lungs: the chemokine (C-C motif) ligand 1 (CCL1), the chemokine (C-C motif) ligand 25 (CCL25), the chemokine (C-X-C motif) ligand 1 (CXCL1), the TNF receptor 1 (TNFR1), interleukin 13 (IL-13) and interleukin 17A (IL17A) (**Fig. 4a** and **b**).

**Figure 4.**
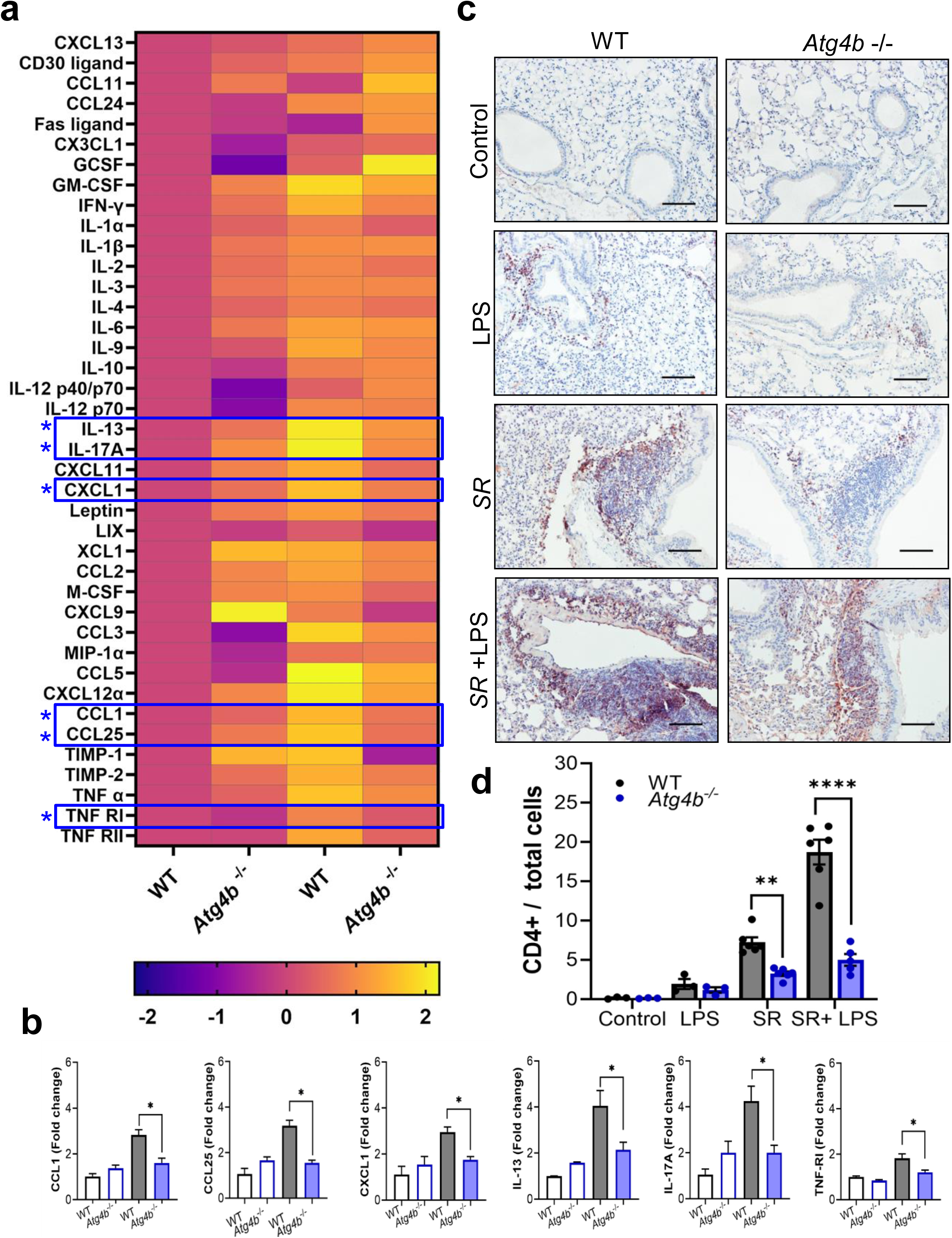
Heat map of multiplex cytokine array from BALFs derived from control and SR+LPS-exposed WT and *Atg4b*^-/-^ mice and CD4+ T cell influx in the lung parenchyma in HP. **a.** Every row represents a protein, and every column represents the mean protein level of each group (Control and SR+LPS-exposed WT and *Atg4b*^-/-^ mice). Fold change was measured relative to the WT control. Data were normalized by log2 transformation. **b**. Bar plots showing the fold change for each upregulated cytokine. **c.** Representative photomicrographs of immunohistochemical positive staining for CD4 in control, LPS-, SR- and SR+LPS-exposed WT and *Atg4b*^-/-^ lungs. Images are at 20X magnification (Scale bars = 100). **d.** CD4-positive cells per total nuclei per experimental group. Data are presented as mean ± SEM. Statistical significance was determined by multifactorial two-way analysis of variance (ANOVA) followed by a Tukey post hoc test (**p<0.005 and ****p<0.005).

CD4+ T cells are the central drivers of HP disease [16, 17]. In this context, we aimed to explore the influx of CD4+ T lymphocytes in *Atg4b*^-/-^ and WT lung parenchyma. We found an increased number of CD4+ T-cells in the intraalveolar spaces, and in the peribronchial and perivascular regions from both *SR*- and *SR*+LPS-exposed WT compared to *Atg4b*^-/-^ mice. We also observed an increased number of CD4+ T-cells into iBALT areas from *SR*+LPS-exposed WT compared to *Atg4b*^-/-^ mice (**Fig. 4c** and **d**). Since we also observed increased levels of the Th2-associated cytokines IL-13 and CCL1 in BALFs from the *SR*+LPS-exposed WT compared to *Atg4b*^-/-^ mice, we evaluated the presence of M2 macrophages within the lung parenchyma. We identified significantly increased YM1 (Chitinase-like 3) (**Fig. 5a** and **b**) and Arginase-1 (ARG1) positive M2-like macrophages in lung parenchyma from both *SR* and *SR*+LPS-exposed WT compared to *Atg4b*^-/-^ exposed mice (**Fig. 5c** and **d**).

**Figure 5.**
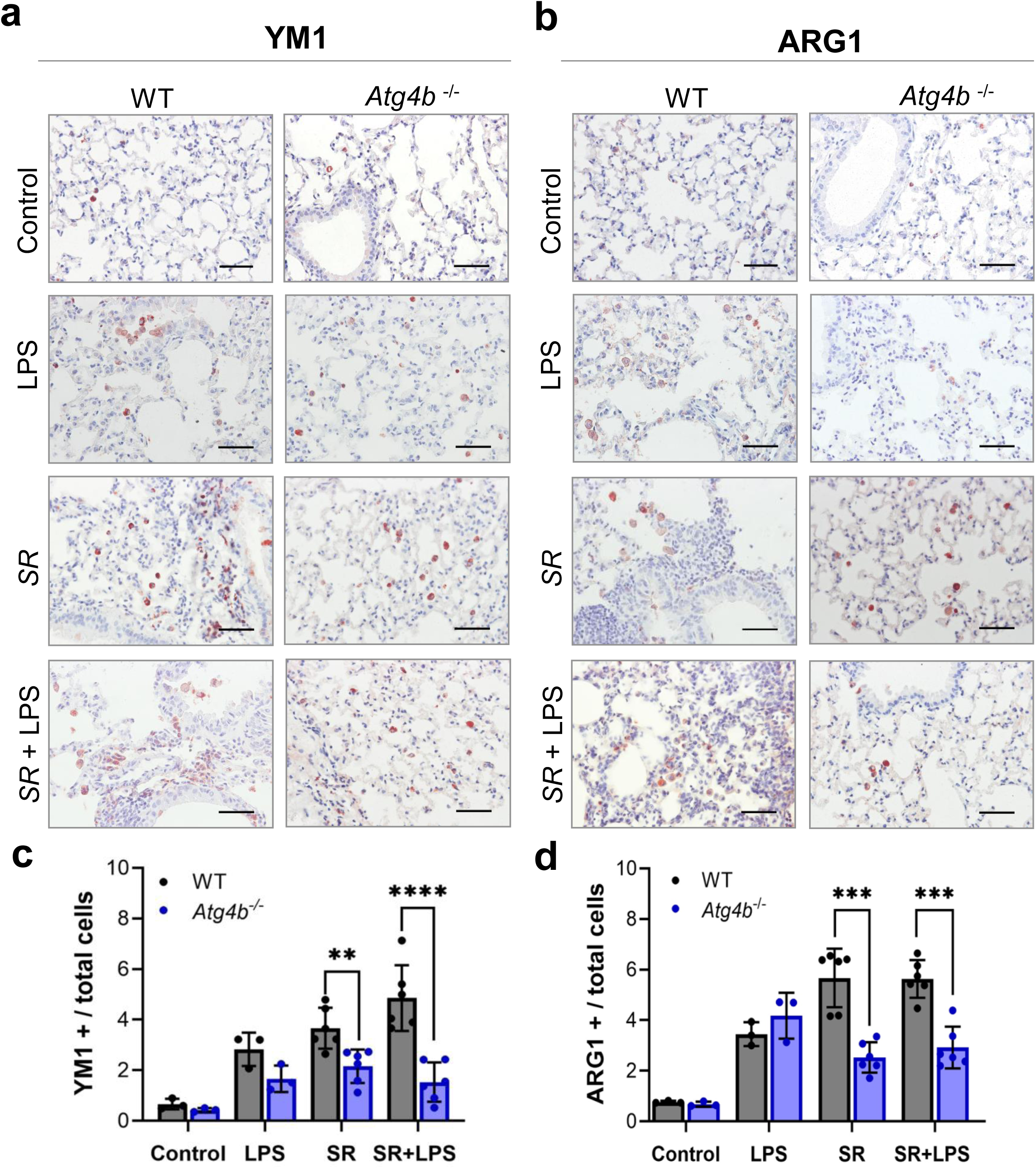
Reduced M2-like macrophages in the lungs from SR- and SR+LPS exposed *Atg4b*^-/-^ mice. **a.** Representative photomicrographs of immunohistochemical positive staining for YM1 and **b.** ARG1 (M2-biomarkers) in control, LPS-, SR- and SR+LPS-exposed WT and *Atg4b*^-/-^ lungs. Positive staining is observed in red, and nuclei are counterstained with hematoxylin in blue. Images are at 40X magnification, and scale bars represent 50 µm. **c.** YM1 and **d.** ARG1 - positive cells per total nuclei per experimental group. Data are presented as mean ± SEM. Statistical significance was determined by multifactorial two-way analysis of variance (ANOVA) followed by a Tukey post hoc test (**p<0.005 and ***p<0.0005).

In summary, our data suggest that autophagy disruption due to the absence of ATG4B results in a reduced inflammatory response associated with a reduced CD4+ lymphocyte infiltration and iBALT area, decreased number of M2-like macrophages, and lower levels of cytokines that perpetuate Th1 and Th17 responses and promote granuloma formation (**Fig. 6a** and **b**). We conclude that autophagy can sustain inflammatory responses during HP pathogenesis, and attenuating this process may ameliorate HP severity.

**Figure 6.**
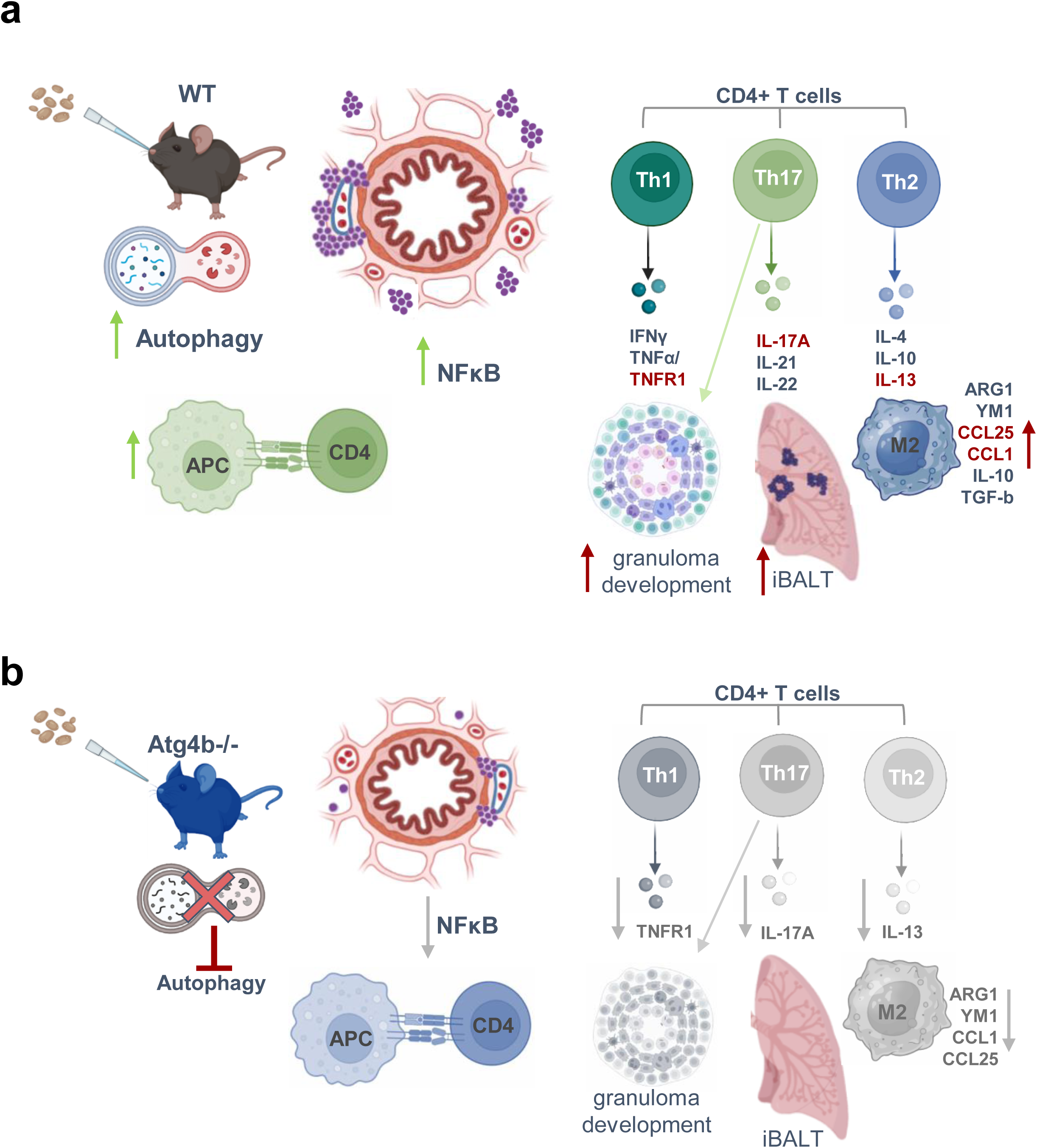
Hypothetical model summarizing the possible mechanisms involved in reduced HP severity in *Atg4b*^-/-^ mice. **a.** Autophagy is induced in macrophages after antigen exposure and is involved in antigen processing and MHC-II presentation and macrophage activation. After TCR engagement, autophagy is involved in CD4+ T cell differentiation and survival. Autophagy modulates NFkB function and cytokine expression. Increased IL-17A leads to granuloma and iBALT development. Increased CCL1 and IL-13 promote M2 polarization. **b.** Loss of autophagy by ATG4B deficiency reduces macrophage activation and NFkB signaling. Autophagy dysfunction also limits CD4+ T cell differentiation and survival, reduces IL-13 and CCL1 signaling that dampens M2 polarization, and reduces IL- 17A signaling that restrains granuloma and iBALT development. This image was created with Biorender.

## DISCUSSION

HP is a dominant immune-mediated ILD, and given that autophagy is a central orchestrator in inflammation, we became interested in studying how autophagy modulates the immune response in this disease [1–5, 18, 19]. Increased LC3B and ATG4B was identified in the bronchial and alveolar epithelium, and in inflammatory cells in the lungs from HP-mouse model, corroborating our previous findings, where several autophagy-related proteins were significantly increased in epithelial and inflammatory cells in lungs from HP patients compared to control subjects [9]. GFP-LC3 puncta colocalization with CD68 in lungs from transgenic mice after *SR* exposure indicates that autophagy activation occurs in macrophages during HP. Macrophages are key antigen-presenting cells that initiate and sustain inflammation in HP [1–5]. Autophagy can regulate the MHC-II antigen-processing pathway and influences both monocyte-to-macrophage differentiation and macrophage polarization [20, 21]. Autophagy activation promotes M2-type polarization and is required for ARG1 expression [22, 23]. Furthermore, the accumulation of M2-polarized macrophages overexpressing MMP14 has been linked to increased severity of *SR*-induced experimental HP, and inhibition of NFκB activation reduced M2-macrophages and HP severity [24]. Autophagy is also required for NFκB activation, since depletion of ATG5, ATG7, Beclin 1, or VPS34 in human cancer cell lines blocked the NFκB signaling pathway [25].

On the other hand, IL-13 and CCL1, two Th2 known drivers of M2-polarization [26], were increased in WT but not in *Atg4b*-deficient HP lungs. CCL1 induced the M2 phenotype in a model of bleomycin-induced fibrosis and is upregulated in the lung tissue of patients with IPF [27, 28]. Consistent with this, reduced CCL1, IL-13 and NFκB levels may contribute to the decreased accumulation of M2-like macrophages in *Atg4b*^⁻/⁻^ HP lungs.

Autophagy is also essential for T cell survival and differentiation [29, 30]. *Atg5* and *Atg*7-deficient mice showed a reduced number of T cells in both the thymus and the periphery, indicating autophagy is necessary for T cell survival [30]. Furthermore, mature peripheral survival is limited in *Beclin1*- or *Vps34*-deficient T-cells, and *PIK3C3-Vps34* deficient CD4+ T cells lose their capacity to differentiate into Th1 subset [31]. In our model, autophagy deficiency induced by *Atg4b* deletion resulted in a decreased number of CD4+ T cells in intraalveolar, peribronchial and perivascular, and iBALT regions, which may contribute to reducing the severity of experimental HP.

Finally, autophagy regulates, and is regulated by several cytokines [32]. Inhibition of autophagy protects against sepsis by attenuating the cytokine storm in a mouse model induced by LPS stimulation and *Escherichia coli* infection [33]. Here, we found a reduced level of CCL1, CCL25, CXCL1, TNFRI, IL-13 and IL17A in the HP lungs from *Atg4b*^-/-^ compared to WT mice. CCL25 regulates T lymphocyte development and tissue-specific homing, and it is among the top 20 upregulated genes in fibrotic HP patients compared to IPF patients and controls [34, 35]. In this context, reduced CCL25 levels could be associated with the reduced number of CD4+ T cells in *Atg4b*^-/-^ HP lungs. However, further mechanistic studies are needed to elucidate the role of CCL1 and CCL25 in HP pathogenesis.

CXCL1 has also been reported to be upregulated in an experimental HP model induced by *Aspergillus niger* plus LPS, and it likely contributes to the amplification of acute inflammatory responses and lung tissue damage [36]. Regarding TNFR-1, the expression of this receptor, along with TNFR-2 and Fas is increased in BALF cells from patients with HP compared to those with IPF and healthy controls [37]. Consistently, *Tnrf1-*deficient mice showed decreased inflammation after *SR* exposure compared to WT [38].

HP was initially considered a Th1-mediated disease, but subsequent studies shown a shift toward a Th17 response after repeated *SR*-exposure [39]. Moreover, while murine HP models incompletely recapitulate the robust granuloma formation observed in human disease, increased IL-17A expression, as found in our study in *SR*+LPS-exposed WT mice, has been shown to promote granuloma development. Supporting this notion, an increased granuloma formation was reported in *T-bet-*deficient mice exposed to *SR* characterized by an exacerbated Th17 cell response, while *Il17ra*-deficient mice were protected against *SR*-induced experimental HP [40, 41]. Moreover, IL-17A/ IL-17Ra pathway contributes to pulmonary iBALT formation. Thus, studies on *Il17a*^−/−^and *Il17ra*^−/−^ mice revealed reduced iBALT formation after *Mycobacterium tuberculosis* infection [42, 43]. Together, this evidence suggests that reduced IL-17A expression in Atg4b-/ lungs could ameliorate granuloma and iBALT development in *Atg4b*^-/-^ mice.

In conclusion, our findings identify autophagy as a critical contributor to HP pathogenesis. Excessive autophagy activation appears to amplify acute inflammatory responses in the lung. Collectively, our results support the concept that modulation of autophagy may represent a promising therapeutic strategy to attenuate disease severity and potentially prevent acute exacerbations and progression to fibrotic HP.

## Declarations

### Author contributions

Conceptualization, funding acquisition and project administration were performed by Sandra Cabrera. Material preparation, data collection and analysis were performed by Andrea Sánchez, Sandra Cabrera, Miguel Gaxiola and Ángeles García. Design of figures was performed by Sandra Cabrera and Andrea Sánchez. The first draft of the manuscript was written by Sandra Cabrera. Manuscript was critically reviewed and edited by Annie Pardo and Moisés Selman. All authors read and approved the final manuscript.

### Ethics approval

Animal research protocols were approved by the Bioethics Committee of the Faculty of Sciences from UNAM (protocol approval CEARC_PI_2022_23_07_Cabrera) and from INER (B24-23) and performed in strict accordance with approved guidelines.

### Data availability

The data generated and analyzed during the current study are available from the corresponding author upon reasonable request.

## Acknowledgments

We thank the Animal Facility staff at INER (Instituto Nacional de Enfermedades Respiratorias “Ismael Cosío Villegas”) for their support and assistance. We thank LANSBIODYT Lab and UNICUA-UNAM Facility, and Dr. Edgar Jiménez-Díaz for the technical support during confocal images capture. We thank Christian Alan Cabello Hernández for their support in developing the HP mouse model.

## SUPPLEMENTARY INFORMATION

**Supplementary figure 1.**
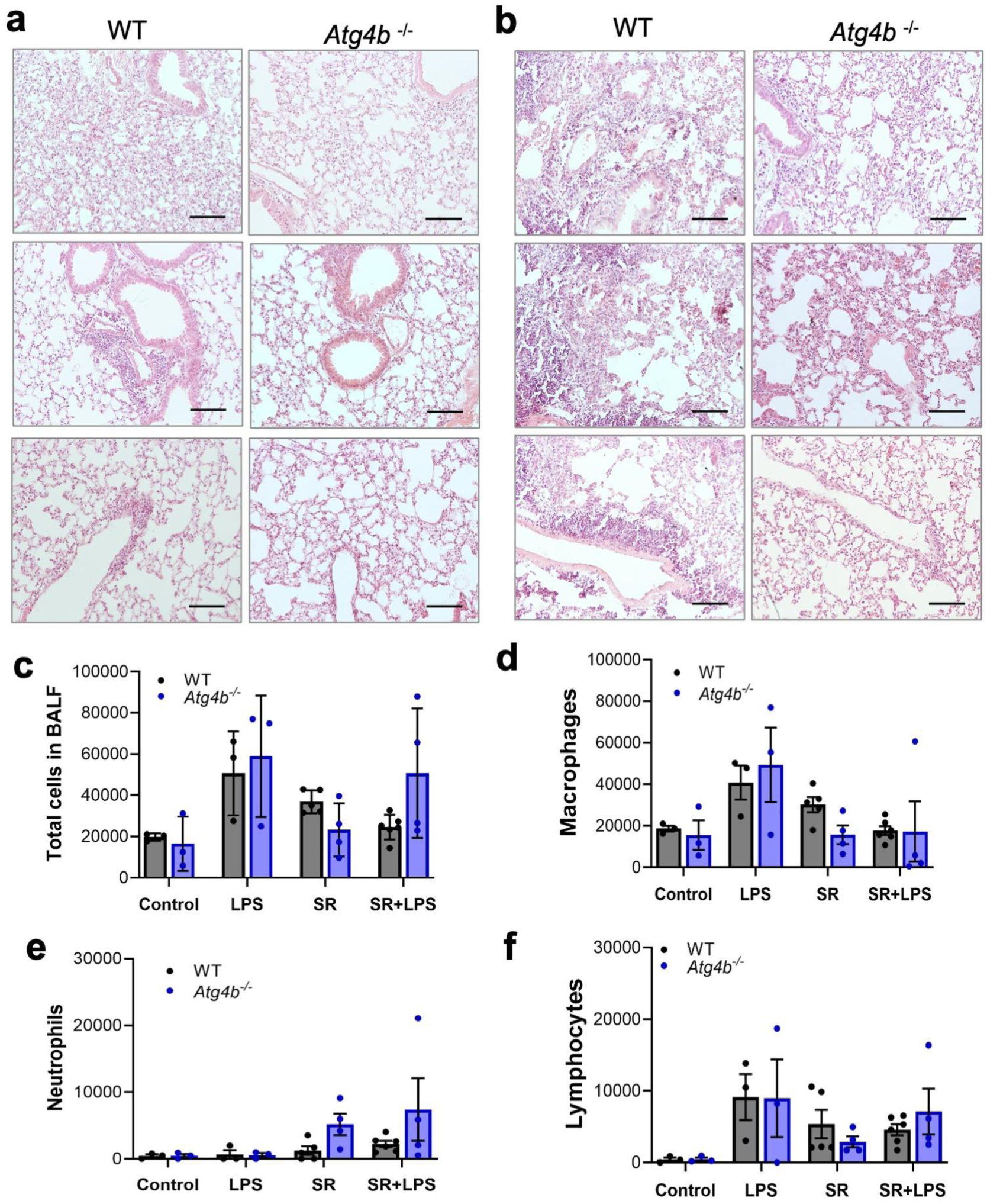
Lung parenchyma from SR and SR+LPS from WT compared to *Atg4b*^-/-^ mice. **a.** Representative light microscopy images of H&E-stained lung tissue sections from SR and **b.** SR+LPS-exposed WT compared to *Atg4b*^-/-^ mice, showing intraalveolar, peribronchial and perivascular inflammation. Images are at 20X, and scale bars represent 100 µm. **c.** Total and differential cell count of **d.** Macrophages, **e.** Neutrophils, and **f.** Lymphocytes in the BALF from control, LPS-, SR- and SR+LPS-exposed WT and *Atg4b*^-/-^ mice. Data are presented as mean ± SEM.

**Supplementary figure 2.**
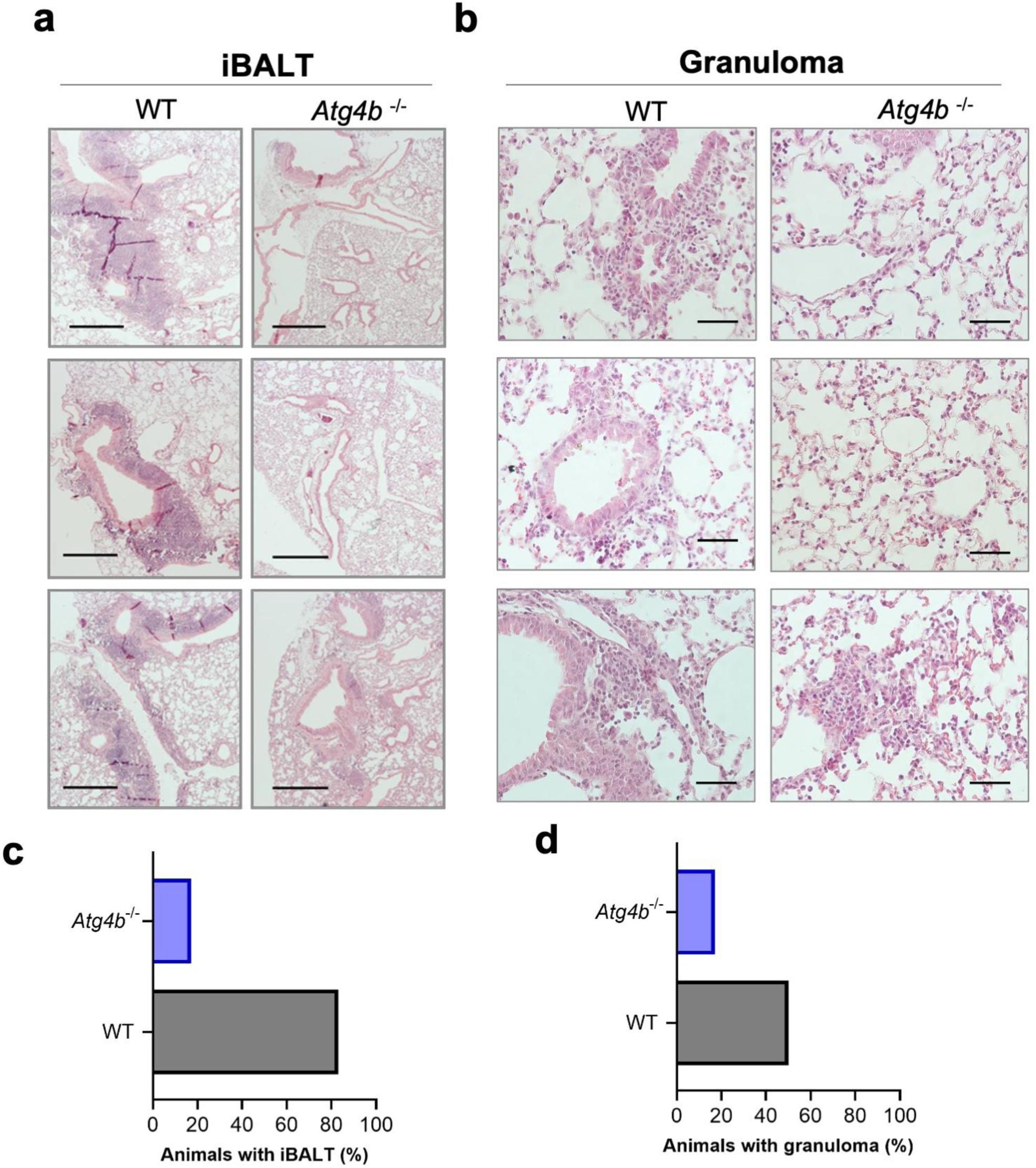
iBALT and granuloma development in SR+LPS exposed WT compared to *Atg4b*^-/-^ mice. **a.** Representative light microscopy images of H&E-stained lung tissue sections from SR+LPS-exposed WT compared to *Atg4b*^-/-^ mice. Images are at 4X, and scale bars represent 250 µm. **b.** Representative light microscopy images showing granulomas in SR+LPS-exposed WT compared to *Atg4b*^-/-^ mice. Images are at 40X, and scale bars represent 50 µm. C. Bars represent the percentage of animals per genotype with iBALT and granuloma development after SR+LPS treatment.

